# Locomotion-induced gain of visual responses cannot explain visuomotor mismatch responses in layer 2/3 of primary visual cortex

**DOI:** 10.1101/2022.02.11.479795

**Authors:** Anna Vasilevskaya, Felix C. Widmer, Georg B. Keller, Rebecca Jordan

## Abstract

The aim of this work is to provide a comment on a recent paper by Muzzu and Saleem (2021). In brief, our concern is that the authors claim that visuomotor mismatch responses in mouse visual cortex can be explained by a locomotion-induced gain of visual halt responses, without directly comparing these responses to mismatch responses. Without a direct comparison, the claim that one response can explain the other appears difficult to uphold, more so because previous work finds that a uniform locomotion-induced gain cannot explain mismatch responses. To support these arguments, we analyzed a series of layer 2/3 calcium imaging datasets and show that coupling between visual flow and locomotion greatly enhances mismatch responses in an experience dependent manner compared to halts in non-coupled visual flow. This is consistent with mismatch responses representing visuomotor prediction errors. Thus, we conclude that feature selectivity cannot explain mismatch responses in mouse visual cortex.

---

Predictive processing is a theoretical framework for a computational description of brain function. It postulates that an internal model of the environment is learned and used to predict sensory inputs (Bastos et al., 2012; Clark, 2013; Jordan and Rumelhart, 1992; Keller and Mrsic-Flogel, 2018; Koster-Hale and Saxe, 2013; Rao and Ballard, 1999). A key neural component of these models is prediction error neurons that compute and report discrepancies between actual sensory input and the prediction of sensory input. Prediction error signals, in turn, are used to drive corrective learning in the internal model. Predictive processing has long been hypothesized as a framework capable of describing cortical function (Rao and Ballard, 1999), but physiological evidence for this has only started to accumulate in the last decade. Much of this evidence in the primary visual cortex (V1) has come from head-fixed mice locomoting in a virtual reality environment with visual feedback coupled to locomotion speed. Pyramidal neurons in layer 2/3 of V1 respond strongly when mismatches between locomotion speed and visual flow speed are presented, which consist of a brief halt in otherwise coupled visual flow during locomotion (Keller et al., 2012). One parsimonious explanation for these responses is that individual neurons compute differences between an inhibitory visual flow driven input and a top-down excitatory prediction of visual flow based on locomotion (Attinger et al., 2017; Jordan and Keller, 2020; Leinweber et al., 2017). This would constitute a negative prediction error computation, driving spiking responses when there is less visual flow than expected based on locomotion. Several key pieces of evidence point toward mismatch responses resulting from this computation, which are summarized in **Figure 1**.

**Figure 1.**
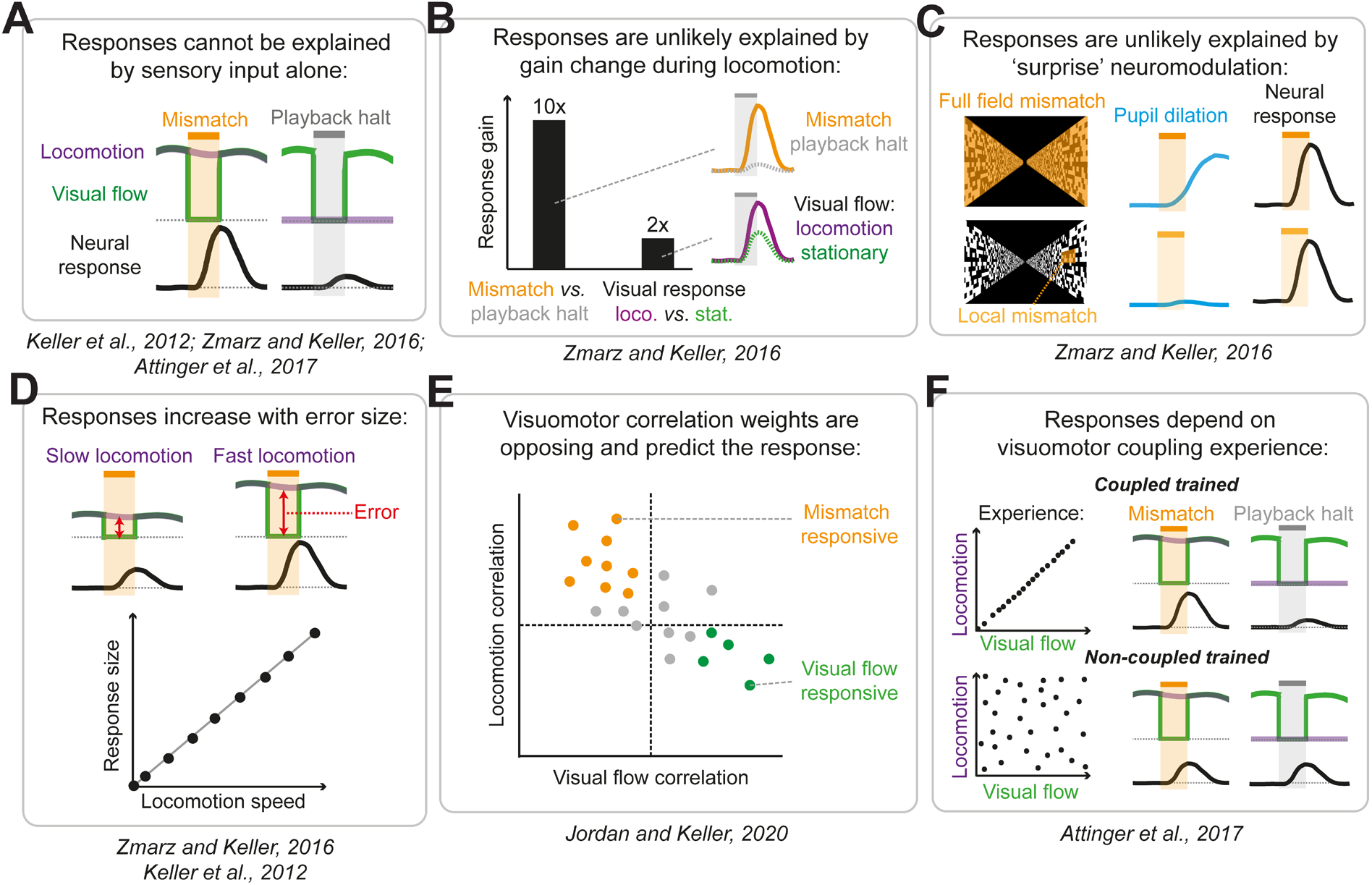
Graphical summary of the key evidence that visuomotor mismatch responses in layer 2/3 of V1 arise from prediction error computation. Prediction error computation consists of a subtractive (or divisive, see (Spratling, 2019)) comparison between sensory input and a prediction of that sensory input, for instance based on movements like locomotion. This can be achieved using a balance of excitation and inhibition from sensory and prediction sources. The key pieces of evidence supporting the idea that visuomotor mismatch responses in layer 2/3 of V1 represent prediction error signals are as follows: (**A**) Mismatch responses cannot be replicated with the visual flow halts alone, uncoupled from locomotion (Attinger et al., 2017; Keller et al., 2012; Zmarz and Keller, 2016). Here and in the subsequent panels of the figure the black lines illustrate the mean population response of pyramidal cells in layer 2/3 of V1. (**B**) The responses cannot be explained by a uniform gain increase of visual responses during locomotion (explained in detail in the main text) (Zmarz and Keller, 2016). The lines on the right illustrate the population responses to the different stimuli (orange = mismatch, gray = playback halt during stationary periods, purple = visual flow during locomotion, green = visual flow during stationary periods). (**C**) Responses likely cannot be explained by a surprise induced neuromodulatory signal, since calcium responses to local mismatches in small parts of the visual field evoke no pupil dilation (which is a proxy for neuromodulatory activity), but are equivalent in size to responses to full field mismatches (which do evoke pupil dilation) (Zmarz and Keller, 2016). (**D**) Responses scale linearly with the degree of error between locomotion speed and visual flow speed (Keller et al., 2012; Zmarz and Keller, 2016). (**E**) Prediction errors should be computed from the difference between visual flow and locomotion speed, and visual flow and locomotion have opposing signs of influence on the membrane potential of layer 2/3 neurons, consistent with a subtractive comparison between the two sources of information (Jordan and Keller, 2020). Somatostatin positive interneurons are consistent with providing visually driven inhibition onto mismatch-responsive neurons (Attinger et al., 2017). (**F**) Mismatch responses depend on the visuomotor coupling experience of the animal: mismatch responses are indistinguishable from passive visual flow halts if the animal is raised with no coupling between visual flow and locomotion (Attinger et al., 2017).

The main claim of the recent paper by Muzzu and Saleem (Muzzu and Saleem, 2021) is that responses to visual flow halts, enhanced by locomotion, could instead account for the visuomotor mismatch responses described in the literature. Please note, we are not questioning the validity of the results presented in the paper, and testing and falsifying models proposed by previous work is absolutely essential. However, with this paper, we would contend that the main conclusion – that visuomotor mismatch responses can be explained by visual flow offset responses modulated by locomotion – is poorly backed up by the data, primarily because the mismatch responses the work claims to be able to explain are never actually measured.

It is important to note that ‘visuomotor mismatch’ describes a stimulus comprising a perturbation (a sudden and brief reduction of visual flow speed to zero) in a condition of closed loop coupling between locomotion and visual flow feedback (see **Figure 1A**). The authors argue that responses to such mismatches can be explained without precise coupling between visual flow and locomotion speeds. Instead, they argue that mismatch responses could result from sensory responses to visual flow halts that undergo a general gain increase during locomotion. The three pieces of evidence used to back up this claim are as follows:

A. Passively presented (open loop) visual flow halts can evoke increased firing in cells that are tuned to low temporal frequencies of drifting visual gratings.
B. These responses can be amplified by locomotion.
C. These responses are not biased toward the forward-backward direction across the population.

Points **A** and **B** have been previously addressed by the visuomotor mismatch literature. More importantly, the experiments in Muzzu and Saleem (2021) do not show that visuomotor mismatch responses are quantifiably explained by these phenomena. Regarding point **A**, it has been demonstrated previously that visual flow halts (termed ‘playback halts’ in the literature) evoke responses in V1 neurons, similar to the stimuli presented in the current paper (Attinger et al., 2017; Keller et al., 2012; Zmarz and Keller, 2016). Regarding point **B**, it has also been shown that locomotion increases the gain of visual responses, on average 2-fold (Bennett et al., 2013; Niell and Stryker, 2008; Polack et al., 2013; Zmarz and Keller, 2016). However, this locomotion-induced gain is on average five-fold smaller than the gain calculated between responses to visuomotor mismatches and playback halts (Zmarz and Keller, 2016). That is – on average, calcium responses to visuomotor mismatch are ten times larger than responses to the identical visual flow stimulus played back to a stationary mouse (playback halts) (**Figure 1B**). This is a key piece of evidence that argues against the idea that locomotion-based amplification of visual responses explains visuomotor mismatch responses. The experiments in Muzzu and Saleem (2021) do not address this, given that there is no quantification of responses to mismatches in yoked visual feedback during locomotion. It is noted in the discussion the possibility that the coupled visual flow condition gives rise to true mismatch responses. However, we are of the opinion that this should be experimentally addressed via quantification of mismatch responses if the conclusion of the paper is that ‘*Feature selectivity can explain mismatch signals in mouse visual cortex*’.

Point **C** is based on the argument that evidence of a V1 population bias for the horizontal direction of visual flow halts (forwards and backwards) is required for visuomotor mismatch signaling. The authors do not find this bias in their recorded population in awake mice. Interestingly, a population bias for naso-temporal visual flow selectivity has been found in the retina, and in layer 2/3 of V1 in anesthetized mice, suggesting it exists in the feedforward signal (Hillier et al., 2017). While it makes sense to match the dynamic range of your sensors to the ethological relevance and statistical properties of the input, we would argue that a bias toward horizontal visual flow preference is not a prediction or a requirement for visuomotor mismatch signaling. Visual responses in V1 have long been known to have orientation/direction tuning, representing the full range across the population. In a freely moving animal, visual flow co-occurs not only with forward locomotion, but with angular velocities of the head in all directions, for instance during turns, rears, and other head movements. These movements will result in visual flow in all directions, and such head movements, in particular turns, will be correlated with locomotion behavior. It would therefore make sense for V1 to have neurons representing visuomotor mismatches with direction tuning covering the full range. The mismatch responses reported in the literature are feature-selective in the spatial domain, with retinotopic receptive fields similar in size to those for classical visual stimuli (Zmarz and Keller, 2016), and it would not be surprising to find that they are also orientation/direction tuned given the known prevalence of such tuning in V1, though this has not yet been directly assessed. Head-fixing an animal and greatly restricting the possible directions of motion to only forwards-backwards should not result in an immediate re-organization of the circuit to overrepresent mismatches along this single axis of motion. In any case, given that the mice in Muzzu and Saleem (Muzzu and Saleem, 2021) never have experience of head-fixed coupled visual flow, and that the stimuli presented are not visuomotor mismatches, we cannot know whether the orientation/direction tuning profile across the population generalizes to mismatch stimuli.

Finally, mismatch responses and evidence of their comparative computation have largely been reported for layer 2/3 in detail (Attinger et al., 2017; Jordan and Keller, 2020; Keller et al., 2012; Zmarz and Keller, 2016), given that deeper layers show few mismatch responses, and show different visuomotor integration characteristics that are not consistent with error-computation (Jordan and Keller, 2020). It is therefore difficult to directly relate the findings of this paper to the known work on mismatch responses in layer 2/3 when it is not reported which layer(s) these unit recordings were made from.

At this point it is important to note, that predictive processing is just one of many possible models that would explain mismatch responses. Muzzu and Saleem’s (Muzzu and Saleem, 2021) claim is that a visual halt response combined with a locomotion-induced gain of visual responses would also explain mismatch responses. As was shown in (Zmarz and Keller, 2016), where the locomotion-induced gain of flow halts was directly compared to the locomotion-induced gain of flow onsets, the effect of locomotion on visual stimuli is not uniform. The apparent gain is substantially larger for flow halts than it is for flow onsets (i.e., classical visual stimuli). Thus, an explanation for mismatch responses based on locomotion related gain would require a visual response to the flow halt combined with a locomotion gain that is stimulus specific.

To directly test whether a stimulus specific locomotion gain could explain layer 2/3 mismatch responses, we compared the locomotion-induced gain of visual flow halt responses in closed and open loop conditions (**Figure 2A**). Assuming a stimulus specific locomotion gain is sufficient to explain mismatch responses, then precise coupling between visual flow and locomotion should have little effect on the size of mismatch responses. In other words, responses evoked by mismatch in the closed loop condition should be very similar to responses evoked by passive visual flow halts during locomotion in the open loop condition. To directly assess this, we analyzed three previously acquired two-photon calcium imaging datasets, recorded from layer 2/3 of V1, totaling 7094 neurons in 45 mice (see Methods). Two of these have been previously published and are publicly available (Attinger et al., 2017; Widmer and Keller, 2021), while one is not yet published. Mice were first exposed to the closed loop condition where visual flow was coupled to locomotion speed, and sudden 1 s halts in visual flow during locomotion were used to evoke mismatch responses. This was followed by an open loop condition where the self-generated visual flow from the preceding closed loop session was replayed to the mouse uncoupled from locomotion. The 1 s halts in visual flow (‘playback halts’) could therefore occur either during stationary periods or during locomotion. Consistent with locomotion increasing the gain of visual responses, in the open loop session we found that playback halts during locomotion evoked a larger population response compared to playback halts during stationary periods (playback halt response, mean ± standard deviation (SD) ΔF/F: stationary: 0.3 ± 5.8 %, locomoting: 1.1 ± 8.7 %, p < 10^-10^, paired t-test). However, mismatches in otherwise coupled visual flow evoked significantly larger responses, on average double the size of those evoked by playback halts during locomotion (mismatch response, mean ± SD ΔF/F: 2.2 ± 9.7 %; mismatch vs playback halt during locomotion, p < 10^-10^, paired t-test) (**Figure 2B and 2E**). These differences cannot be accounted for by differences in locomotion speed, as average locomotion speeds in both conditions were similar (**Figure 2C**). By experimental design, the only difference between the two conditions was the coupling between visual flow and locomotion preceding the visual flow halt (**Figure 2D**).

**Figure 2.**
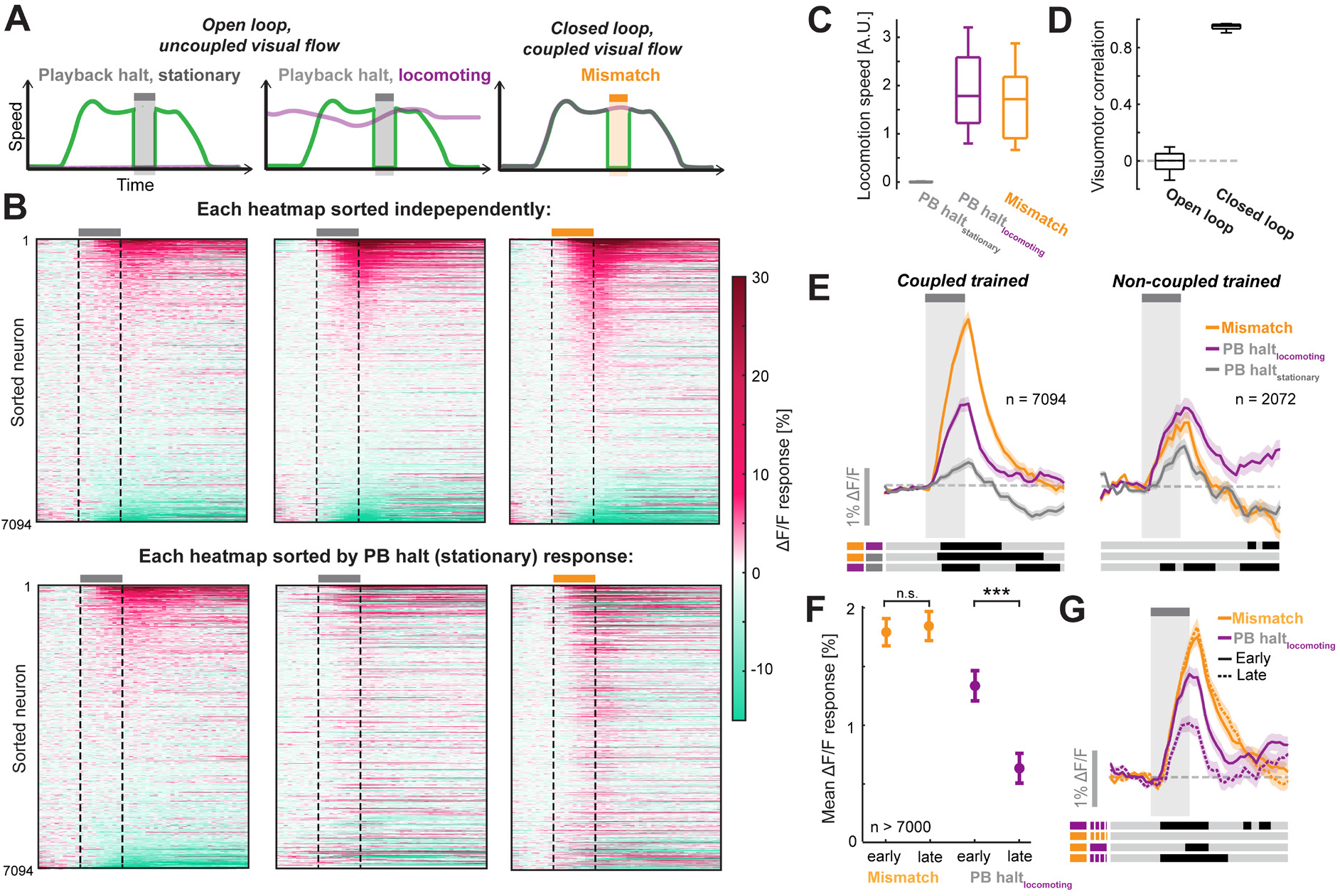
Visuomotor coupling enhanced mismatch responses in an experience dependent manner. (**A**) Schematics to show the three conditions in which responses to visual flow halts were assessed. The green line illustrates visual flow speed, and the purple line illustrates locomotion speed. Left: Mice are stationary while observing a visual flow halt (playback halt, gray shading). Middle: Mice are locomoting and observe a visual flow halt, but locomotion and visual flow are not coupled. Right: Mice observe a visual flow halt while locomoting in a closed loop condition. These events we refer to as visuomotor mismatches (orange shading). (**B**) Heatmaps of the average responses to visual flow halts corresponding to the three conditions in **A**, sorted across 7094 neurons. Top row: Heatmaps were sorted independently for each condition. Bottom row: The same data but with heatmaps that were sorted according to the responses to playback halt, stationary (leftmost heatmap). Note, the correlations between responses in the different conditions were relatively low. (**C**) Average locomotion speed during the visual flow halt stimuli in each condition. (**D**) The correlation between locomotion speed and visual flow speed across the whole session was close to 0 in the open loop condition and close to 1 in the closed loop condition. Note, the only reason closed loop correlation is less than 1 is due to the presentation of mismatches. (**E**) Population average responses to visual flow halt stimuli in the three conditions for mice raised with (left, coupled trained) and without (right, non-coupled trained) visuomotor coupling experience. Shading shows standard error of the mean. Lines below the plots indicate statistical differences between responses as color coded to the left (gray: not significant; black: p<0.01; paired t-test). The comparison being made for each line is indicated by the combination of colors on the left. (**F**) Mean responses to mismatch and playback halt during locomoting in coupled trained animals, in the early and late blocks (see Methods) of closed loop and open loop sessions (where at least 7000 neurons met the criterion of having at least three trials in a given condition). In the experimental paradigm open loop sessions always were always presented after closed loop sessions. Thus, the longer the animal was exposed to an open loop condition, the smaller the playback halt locomoting responses became. n.s.: p > 0.05, ***: p < 0.001, paired t-test. (**G**) Population average responses to mismatch (orange) and playback halts during locomotion (purple) in coupled trained animals, in the early and late blocks (see Methods) of closed loop and open loop sessions respectively (where at least 7000 neurons met the criterion of having at least three trials in a given condition). Shading shows standard error of the mean. Lines below the plots indicate statistical differences between responses as color coded to the left (gray: not significant; black: p<0.01; paired t-test). The comparison being made for each line is indicated by the combination of colors on the left.

This result is consistent with the idea that in the closed loop condition, strong predictions of visual flow during locomotion drive prediction error responses when the visual flow is lower than expected, and directly opposes the idea that a uniform locomotion-induced gain of visual flow halt responses, whether stimulus specific or not, can explain mismatch responses. Note that predictions likely still play a reduced role in the enhancement of playback halt responses during locomotion in the open loop condition, given that mice have a lifetime of experience with coupled locomotion and visual flow. Thus, the effect of visuomotor coupling on mismatch responses shown here is potentially an underestimate. To assess locomotion related gain in mice raised without coupling between forward locomotion and backward visual flow (non-coupled trained), we assessed visual flow halt responses in mice raised in an open loop virtual environment (2072 neurons from 9 mice (Attinger et al., 2017)). In these mice, locomotion increased playback halt responses (playback halt response, mean ± SD ΔF/F: stationary: 0.4 ± 5.0 %, locomoting: 1.2 ± 6.4 %, p < 10^-7^, paired t-test), but the enhancing effect of visuomotor coupling on mismatch responses was entirely absent (mismatch response, mean ± SD ΔF/F: 0.9 ± 6.7 %; mismatch vs playback halt during locomotion, p < 0.05, paired t-test) (**Figure 2E**).

An interesting question the results of Muzzu and Saleem raise is, on what timescale does the experience of coupling, or the absence of coupling, shape visuomotor prediction error responses? Exposed to an open loop environment, the system likely gradually reduces predictions of visual flow given movement, thereby decreasing prediction error responses. If mice are raised without visuomotor coupling, mismatch responses are not increased by visuomotor coupling, consistent with a lack of visuomotor prediction error responses (**Figure 2E**). When transitioning from a closed loop virtual environment to an open loop virtual environment, locomotion onset related mismatch responses decay with a half-life of a few minutes in normally raised animals (see Figure 3B in Keller et al., 2012). To test whether playback halt responses during locomotion undergo a similar reduction in amplitude as a function of time since the start of the open loop condition, we compared these responses between early (0 s to 250 s) and late (500 s to 750 s) segments of the open loop condition in mice raised with visuomotor coupling. All experiments started with a closed loop condition that was followed by an open loop condition. In mice raised with visuomotor coupling, the size of playback halt responses during locomotion declined between the early and late periods in the open loop session, toward a value close to the average playback halt response during stationary periods (playback halt response during locomotion, mean ± SD ΔF/F: early: 1.4 ± 11.3 %, late: 0.7 ± 10.7 %, p < 10^-10^, paired t-test) (**Figure 2F-G**). This reduction in response could not be explained by a general time-dependent change, since mismatch responses showed no sign of reduction between early and late epochs of the closed loop session (mismatch response, mean ± SD ΔF/F: early: 1.8 ± 11.1 %, late: 1.9 ± 11.9 %, p = 0.90, paired t-test). Neither could the reduction in response be explained by a reduction in locomotion speed (p = 0.92, paired t-test between locomotion speeds in early and late epochs). These results are consistent with locomotion-induced gain being strongly dependent on the recent experience of visuomotor coupling.

Thus, visuomotor mismatch responses can be described either as a stimulus specific and experience dependent locomotion gain, or as a visuomotor prediction error. Both are technically valid explanations of mismatch responses. However, we would argue that a stimulus specific and experience dependent locomotion gain is not a useful model in the sense that it makes experimental predictions. It is a list of observations that does not address the question of why it would make sense to compute such a signal. Most importantly such a model is not normative and fails to predict the results summarized in **Figure 1**. By contrast, a predictive processing model that interprets mismatch responses as prediction errors explains all of these results and has direct computational application (Lotter et al., 2016; Straka et al., 2020; Whittington and Bogacz, 2017).

In summary, our primary concern with the alternative explanation for visuomotor mismatch responses in V1 provided by Muzzu and Saleem (2021) is that visual flow is never coupled to locomotion, and therefore mismatch responses are not quantified. We argue that without such quantification it is difficult to provide evidence for a useful alternative explanation for mismatch responses, or any other neural response. The comparison between responses to passive visual flow halts and mismatches we present here (**Figure 2**) provides this quantification and, alongside previous findings (**Figure 1**), demonstrates that locomotion-induced gain cannot account for mismatch responses in layer 2/3 of V1. Overall, we consider the results presented in Muzzu and Saleem (2021) both valid and valuable for the field, however, we find the interpretation and the title misleading due to the concerns we outlined here.

## METHODS

All details of data acquisition were described previously (Attinger et al., 2017; Widmer and Keller, 2021). Data are publicly available on https://data.fmi.ch/PublicationSupplementRepo/.

### Description of the dataset

Raw data consisted of two-photon calcium imaging recordings from V1 layer 2/3 pyramidal neurons, totaling 10005 neurons recorded in 66 mice with visuomotor experience, and 2561 neurons recorded in 9 mice raised with only open loop visual experience. Neurons undergoing analyses were selected based on criteria below, described under ‘Calculation of responses and exclusion criteria’). Data were recorded at 10 Hz or 15 Hz. 15 Hz data was down sampled to 10 Hz for analysis. All data comprised the following stimulus presentation structure. First, a session with 500 s or 750 s closed loop virtual reality experience, in which visual flow was yoked with locomotion speed. In this condition, brief 1 s visual flow halts were presented to the mice during locomotion, which we refer to as mismatch stimuli. Next, two or three open loop sessions were presented with the same duration as the closed loop session, in which visual flow recorded from the initial closed loop sessions was replayed to the mouse, and thus not yoked to locomotion. In this condition, the visual flow halts were referred to as playback halts and could occur both during locomotion and stationary behavior.

### Definition of locomotion and stationary trials

Locomotion speed for each trial was assessed by averaging the locomotion speed in a time window of −1.5 to +1.5 s from stimulus onset. Due to differences in the measurement of locomotion speed across the datasets, the threshold for locomotion was determined as a velocity above 12% of the 95^th^ percentile of the speed distribution within a dataset, while the threshold for stationary periods was defined as a velocity below the 2% of the 95^th^ percentile.

### Calculation of responses and exclusion criteria

For each neuron, the ΔF/F response in each trial was first baseline subtracted using the mean ΔF/F in the 1 s preceding the stimulus, before being averaged across trials. By design, mismatch triggers and playback halts while locomoting occur during locomotion. Hence the probability of locomotion at the time of the trigger is 100% but reduces with increasing time from the trigger. As a result of this, neurons whose activity correlates positively with locomotion will exhibit increases of activity around time of the trigger. To correct for these non-specific increases in ΔF/F during mismatch and playback halts while locomoting we calculated sham responses and subtracted these from mismatch responses and playback halt responses during locomotion. Sham responses were calculated based on 1000 random triggers, selected based on the same locomotion criteria used to calculate actual responses. For **Figure 2B, E**, only neurons with at least 5 trials in all three stimulus conditions (mismatch, playback halt during locomotion and during stationary periods) were included in the analyses (coupled trained, n = 7094; non-coupled trained, n = 2072). To calculate average responses for each neuron (e.g., numbers reported in the text, and in **Figure 2F**), we took the mean of the average response in a window 500 ms to 1500 ms after stimulus onset. In all statistical tests, responses were compared with a paired t-test.

### Early vs late response comparisons

Early vs late response comparisons: To compare responses in early and late parts of the session (**Figure 2F-G**), we took responses during the first 250 s of the first open loop or closed loop sessions to calculate early responses. To calculate late responses, we used responses in the last 250 s of the closed loop condition to calculate mismatches (250 s to 500 s for 3625 neurons with 500 s duration closed loop sessions, or 500 s to 750 s for 5539 neurons with 750 s duration closed loop sessions), while we took the third block of 250 s in the open loop condition (500 s to 750 s) to calculate playback halt responses. To calculate the average responses, only neurons with at least three trials in a given condition (mismatch, playback halt during locomotion, early and late) were included (at least 7000 neurons in each condition).

## ACKNOWLEDGEMENTS

We thank the members of the Keller lab for discussion and support. This work has received funding from the Swiss National Science Foundation, the Novartis Research Foundation, the Human Frontier Science Program (LT000077/2019-L) to R.J., and the European Research Council (ERC) under the European Union’s Horizon 2020 research and innovation programme (grant agreement No 865617).

## AUTHOR CONTRIBUTIONS

RJ performed all analyses, FCW performed the unpublished experiments, AV, GBK, and RJ wrote the manuscript.

## CORRECTION HISTORY

**25.04.2022:** We corrected two errors in our analysis code. These resulted in small changes to Figure 2 and to some of the numbers reported in the manuscript (small changes in number of neurons, and response amplitudes, none of which change the reported comparisons). The cause for the error was that in a subset of imaging sites, closed-loop sessions were longer than assumed in the code and thus some mismatch trials were misclassified as early playback halt trials in Figures 2F-G. Correcting this error did not affect the outcome of any of the comparisons made.

## REFERENCES

Attinger, A., Wang, B., and Keller, G.B. (2017). Visuomotor Coupling Shapes the Functional Development of Mouse Visual Cortex. Cell 169, 1291–1302.e14.

Bastos, A.M., Usrey, W.M., Adams, R.A., Mangun, G.R., Fries, P., and Friston, K.J. (2012). Canonical microcircuits for predictive coding. Neuron 76, 695–711.

Bennett, C., Arroyo, S., and Hestrin, S. (2013). Subthreshold mechanisms underlying state-dependent modulation of visual responses. Neuron 80, 350–357.

Clark, A. (2013). Whatever next? Predictive brains, situated agents, and the future of cognitive science. The Behavioral and Brain Sciences 36, 181–204.

Hillier, D., Fiscella, M., Drinnenberg, A., Trenholm, S., Rompani, S.B., Raics, Z., Katona, G., Juettner, J., Hierlemann, A., Rozsa, B., et al. (2017). Causal evidence for retina dependent and independent visual motion computations in mouse cortex. Nat Neurosci 20, 960–968.

Jordan, M.I., and Rumelhart, D.E. (1992). Forward Models: Supervised Learning with a Distal Teacher. Cognitive Science 16, 307–354.

Jordan, R., and Keller, G.B. (2020). Opposing Influence of Top-down and Bottom-up Input on Excitatory Layer 2/3 Neurons in Mouse Primary Visual Cortex. Neuron 108, 1194–1206.e5.

Keller, G.B., and Mrsic-Flogel, T.D. (2018). Predictive Processing: A Canonical Cortical Computation. Neuron 100, 424–435.

Keller, G.B., Bonhoeffer, T., and Hübener, M. (2012). Sensorimotor mismatch signals in primary visual cortex of the behaving mouse. Neuron 74, 809–815.

Koster-Hale, J., and Saxe, R. (2013). Theory of mind: a neural prediction problem. Neuron 79, 836–848.

Leinweber, M., Ward, D.R., Sobczak, J.M., Attinger, A., and Keller, G.B. (2017). A Sensorimotor Circuit in Mouse Cortex for Visual Flow Predictions. Neuron 95, 1420–1432.e5.

Lotter, W., Kreiman, G., and Cox, D. (2016). Deep Predictive Coding Networks for Video Prediction and Unsupervised Learning. 5th International Conference on Learning Representations, ICLR 2017 - Conference Track Proceedings.

Muzzu, T., and Saleem, A.B. (2021). Feature selectivity can explain mismatch signals in mouse visual cortex. Cell Rep 37, 109772.

Niell, C.M., and Stryker, M.P. (2008). Highly selective receptive fields in mouse visual cortex. The Journal of Neuroscience 28, 7520–7536.

Polack, P.-O., Friedman, J., and Golshani, P. (2013). Cellular mechanisms of brain-state-dependent gain modulation in visual cortex. Nat Neurosci 16, 1331–1339.

Rao, R.P., and Ballard, D.H. (1999). Predictive coding in the visual cortex: a functional interpretation of some extra-classical receptive-field effects. Nat. Neurosci. 2, 79–87.

Spratling, M.W. (2019). Fitting predictive coding to the neurophysiological data. Brain Research 1720, 146313.

Straka, Z., Svoboda, T., and Hoffmann, M. (2020). PreCNet: Next Frame Video Prediction Based on Predictive Coding. ArXiv.

Whittington, J.C.R., and Bogacz, R. (2017). An approximation of the error backpropagation algorithm in a predictive coding network with local hebbian synaptic plasticity (MIT Press Journals).

Widmer, F.C., and Keller, G.B. (2021). Developmental plasticity in visual cortex is necessary for normal visuomotor integration and visuomotor skill learning. BioRxiv 2021.06.20.449148.

Zmarz, P., and Keller, G.B. (2016). Mismatch Receptive Fields in Mouse Visual Cortex. Neuron 92, 766–772.

